# Correlated individual differences suggest a common mechanism underlying metacognition in visual perception and visual short-term memory

**DOI:** 10.1101/140558

**Authors:** Jason Samaha, Bradley R. Postle

**Affiliations:** University of Wisconsin-Madison, Department of Psychology; University of Wisconsin-Madison, Department of Psychiatry

**Keywords:** Metacognition, Visual perception, Short-term memory, Confidence, Signal detection theory

## Abstract

Adaptive behavior depends on the ability to accurately introspect about one’s own performance. Whether this metacognitive ability is supported by the same mechanisms across different tasks has thus far been investigated with a focus on correlating metacognitive accuracy between perception and long-term memory paradigms. Here, we investigated the relationship between metacognition of visual perception and metacognition of visual short-term memory (VSTM), a cognitive function thought to be more intimately related to visual processing. Experiments 1 and 2 required subjects to estimate the perceived or remembered orientation of a grating stimulus and rate their confidence. We observed strong positive correlations between individual differences in metacognitive accuracy between the two tasks. This relationship was not accounted for by individual differences in task performance or average confidence, and was present across two different metrics of metacognition and in both experiments. A model-based analysis of data from a third experiment showed that a cross-domain correlation only emerged when both tasks shared the same task-relevant stimulus feature. That is, metacognition for perception and VSTM were correlated when both tasks required orientation judgments, but not when the perceptual task was switched to require contrast judgments. In contrast to previous results comparing perception and long-term memory, which have largely provided evidence for domain-specific metacognitive processes, the current findings suggest that metacognition of visual perception and VSTM is supported by a domain-general metacognitive architecture, but only when both domains share the same task-relevant stimulus feature.

## Introduction

When humans make decisions they are capable of estimating the likelihood that their decision was accurate. This introspective ability falls under a class of cognitive processes known as metacognition because it entails cognizing about the quality of a decision-making process (1). Intuitively, an individual has high metacognitive accuracy if their estimate of the accuracy of their decision (e.g., as expressed by a confidence rating) corresponds well with the actual accuracy of their decision (2). Because decisions can be made on the basis of information from a plethora of sources—for example, deciding on the basis of current sensory input versus deciding on the basis of information culled from long-term memory—an outstanding question is whether metacognitive processes are domain-general or domain-specific (3). A domain-general metacognitive monitoring process would be expected to evaluate the accuracy of decisions made from both perceptual inputs as well as those based on memory. In contrast, a domain-specific metacognitive system would use independent neural resources or computations to estimate the quality of memory-versus perception-based judgments, for example.

Recent work on this topic has focused on correlating individual differences in metacognition during perception and long-term memory and has resulted in mixed findings. Several studies have reported non-significant relationships between individual’s metacognitive ability in a perceptual task and their metacognitive ability in a long-term memory task (4–6), suggesting domain-specific metacognition. However, an experiment using similar tasks did find a reliable positive correlation between metacognitive abilities in both domains (7), and other work has shown correlated metacognitive performance across different perceptual tasks (8), suggesting some shared underlying resources. A number of the above-mentioned studies, however, have also reported that structural and function brain imaging data from distinct regions correlated with metacognitive abilities for the distinct tasks, reinforcing domain-specificity at the neural level (4,5,7). Additional evidence for domain-specificity between perception and long-term memory has come from a recent study of patients with lesions to anterior portions of prefrontal cortex. These patients showed a selective deficit in visual perceptual metacognition, but not memory metacognition for a recently studied word list (9).

A lack of cross-task correlation in metacognition may sometimes be difficult to interpret because this could result from procedural differences between tasks not necessarily related to the cognitive construct under investigation (e.g., the use of different stimuli in the perception versus memory task). Furthermore, perception and long-term memory are themselves quite distinct cognitive functions (although they can certainly interact in some situations, e.g., (10)), and an underexplored question is whether perceptual metacognition relates to metacognition for other cognitive functions more closely related to perception. Across three experiments, we examined whether metacognition in visual perceptual judgments is related to metacognition for visual short-term memory (VSTM) judgments using tasks with the same stimuli that differ only in the requirement for memory storage over a short delay (Experiments 1 and 2), or tasks that differ also in the relevant stimulus feature (Experiment 3). Because perception of and VSTM for a given stimulus feature are hypothesized to rely on shared neural representations (11–14), we might anticipate that metacognition in these domains is also based on some shared resource, leading to positively correlated individual differences in metacognition across tasks, but only when the task-relevant stimulus feature is shared.

## Materials and Methods

### Data availability

In accordance with the practices of open science and reproducibility, all raw data and code used in the present analyses are freely available through the Open Science Framework (https://osf.io/py38c/).

### Experiments 1 and 2

Because of their similarities, the methods pertaining to Experiment 1 and 2 are described together in this section, followed by the methods for Experiment 3.

### Participants

Forty subjects (twenty in Experiment 1: mean age = 21 years, *SD =* 1.67, 10 female, and twenty in Experiment 2: mean age = 20.6 years, *SD* = 2.01, 14 female) from the University of Wisconsin-Madison community participated in these experiments and received monetary compensation. All subjects provided written consent, reported normal or corrected-to-normal visual acuity and color vision, and were naive to the hypothesis of the experiment. The University of Wisconsin-Madison Institutional Review Board approved all experiments reported here.

### Stimuli

Target stimuli were identical for both experiments and consisted of sinusoidal luminance gratings embedded in white noise presented within a central circular aperture (see Figure 1A). Gratings subtended 2 degrees of visual angle (DVA), had a spatial frequency of 1.5 cycles/DVA and a phase of zero. Fixation (a light gray point, 0.08 DVA) was centered on the screen and was dimmed slightly to indicate trial start (see Figure 1A). Noise consisted of white noise luminance values generated randomly on each trial for each pixel in the noise patch. The contrast of the grating was determined for each subject by an adaptive staircase procedure prior to the main tasks. On a random half of the trials the contrast of both the signal and the noise was halved. This was not expected to impact orientation estimation performance because the signal-to-noise ratio of the stimulus was unchanged (15), however it led to a relatively small but reliable performance difference in Experiment 1 (difference in error = 1.7°, p<0.001), but not in Experiment 2 (difference = 0.15°, p = 0.79). This manipulation was not further explored here. Stimuli were presented on an iMac computer screen (52 cm wide × 32.5 cm tall; 1920 × 1200 resolution; 60 Hz refresh rate). Subjects viewed the screen from a chin rest at a distance of 62 cm. Stimuli were generated and presented using the MGL toolbox (http://gru.stanford.edu) running in MATLAB 2015b (MathWorks, Natick, MA, USA).

**Figure 1.**
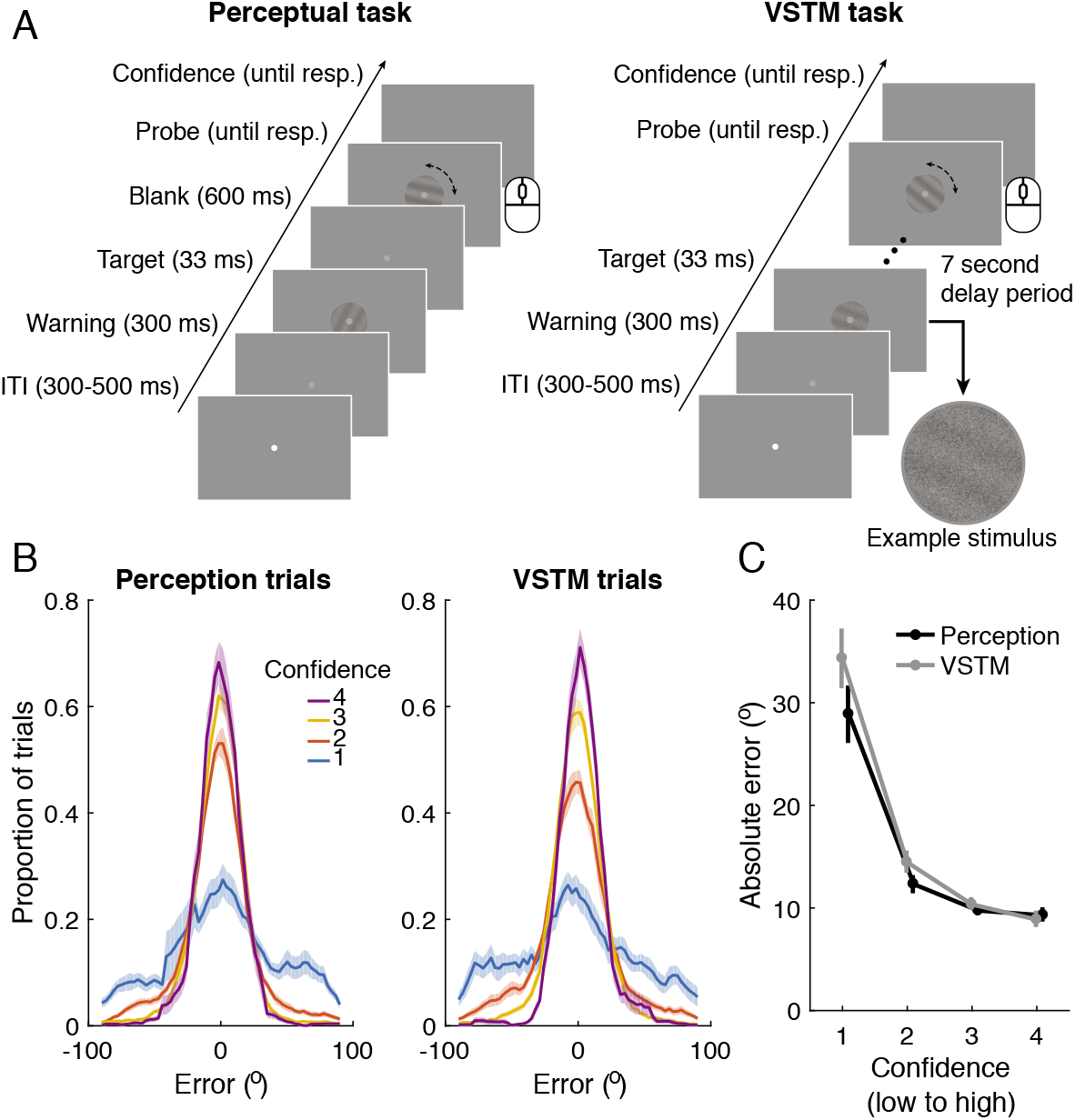
Orientation estimation tasks and confidence-error relationships for Experiment 1. (A) On each trial of the perception and VSTM task, subjects moved a computer mouse to match the perceived or remembered orientation and then provided a confidence rating on a 1-4 scale to indicate how close they thought they came to the true orientation where 1 = “complete guess”and 4 = “very close”. The tasks differed only by the addition of a 7 second delay period for theVSTM task. (B) Distributions of responses relative to the true orientation (i.e., error) show aclear scaling with confidence ratings, suggesting that subjective ratings track objectiveperformance at the group level. (C) Median absolute error scales with confidence and VSTMtrials produced overall greater error, indicating that representations became noisier when heldin VSTM. ITI: inter-trial interval. Shaded bands and error bars denote ± 1 SEM.

### Perceptual task

To probe each individual’s perceptual metacognitive abilities, we employed an orientation estimation task with confidence ratings (16). On each trial, a target grating was presented centrally for 33 ms with a randomly determined orientation between 1-180°, followed shortly (600 ms) by a highly visible probe grating without noise, whose orientation could be rotated via mouse movement. This short interval between the target and probe was necessary to ensure that the probe had no visual masking effect. Subjects were instructed to match the perceived orientation as closely as possible. Subjects pressed the spacebar to input their orientation response and then used number keys 1-4 to provide a confidence rating. Because performance in this task varies continuously (as opposed to a binary correct/incorrect outcome) we instructed subjects to use the confidence scale to indicate how close they think they came to the true orientation using the scale labels 1 = “complete guess” and 4 = “very close”. These perceptual task parameters were the same for both experiments. See Figure 1A for complete trial timings.

### VSTM task

To probe metacognitive abilities for VSTM, we introduced a delay period between the target and the response probe. In Experiment 1, the delay period was fixed at 7 seconds and in Experiment 2 it was randomly sampled from the set: 3.45, 6.30, 9.15, or 12.00 seconds. The stimuli and all other task events were identical to the perceptual task in order to minimize any differences between tasks that are unrelated to the cognitive manipulation of interest

### Staircase

To minimize performance differences across subjects, both experiments began with 100 trials of a 1-up, 3-down adaptive staircase procedure, which classified responses as correct or incorrect depending on whether they were within 25° of a trial’s true orientation. This procedure aimed to produce ∼80% of trials with less than 25° error. The staircase began with the grating component of the stimulus having a Michelson contrast of 50%, which was then averaged with a 100% contrast white noise patch. The step size in grating contrast was adapted according to the PEST algorithm (17), with an initial starting step size of 20% contrast. Procedurally, the staircase task was identical to the perceptual task (described above). The resulting mean (SEM) contrast of the grating (prior to averaging with noise) was 7.8% (0.47) for Experiment 1 and 8.5% (0.61) for Experiment 2, and was held constant throughout the rest of both experiments.

### Procedure

For Experiment 1, perceptual and working memory tasks were performed in separate blocks. Following the staircase, each subject completed one block of 120 trials of the perceptual task, followed by three blocks of 60 trials each of the VSTM task, followed by another block of the perceptual task. This resulted in a total of 240 perceptual trials and 180 VSTM trials per subject, completed in a single 1.5 hour session. Experiment 2 differed in that perceptual and VSTM trials were intermixed within blocks and randomly determined with equal probability to be either a perceptual trial or one of the four delay periods (between 3.45 −12 seconds) of the VSTM task. Intermixing perception and VSTM trials further minimized procedural differences between tasks by eliminating any task-related expecations (since subjects did not know which type of trial would come next) and by removing temporal delays between task performance. Each subject completed 300 trials, seperated into 5 blocks, resulting in an average (± SD) of 55.5 (6.4) perceptual trials and 59 (8.0) trials of each delay period of the VSTM task, after removal of trials based on response times (see below). Total task time was ∼1.5 hours.

### Quantifying metacognition

Task performance is measured as error (in degrees) between the subject’s response and the true orientation on each trial (see Figure 1B). To relate this continuously varying performance metric to subjective confidence ratings, we computed rank correlations between each trials’ absolute error and confidence rating to capture how well confidence tracks performance. Error should decrease with increasing confidence so a subject with good metacognition would have a stronger negative correlation between confidence and error than a subject with poor metacognition. Although intuitive, and used elsewhere (18,19), this metric is potentially influenced by factors not necessarily related to metacognitive accuracy per se, such as task difficulty and biases in confidence scale use (e.g., under or overconfidence; (2)). Although we used a staircase procedure to match difficulty, there was still considerable variability across subjects in median absolute error in both Experiment 1 (range: 8 −16.5°) and Experiment 2 (range: 6.9 −23.3°). A recently introduced measure called meta-d’/d’ can correct for these influences (20), however, meta-d’/d’ has been developed only for tasks with discreet outcomes amenable to signal detection theory analysis (e.g., hits, misses) and cannot be applied to the continuous estimations tasks we employed in Experiments 1 and 2 (but see Experiment 3). In order to control for these influences when testing our primary hypothesis about the relationship between perceptual and VSTM metacognition, we ran two additional multiple regression models that included covariates for average and task-specific error and confidence (see *Statistics* below). In the case of models with multiple predictors, the relationship between perceptual and memory metacognition was visualized (Figure 2 and 4) using added variable plots (MATLAB function *plotAddedm*), which use the Frisch-Waugh-Lovell theorem to partial out the effects of other predictors in the model, revealing the effect of a single predictor while all other predictors are held constant. Predictor R^2^ for these models was computed as the sum of squares for the perceptual metacognition predictor divided by the total sum of squares for all other predictors and error.

**Figure 2.**
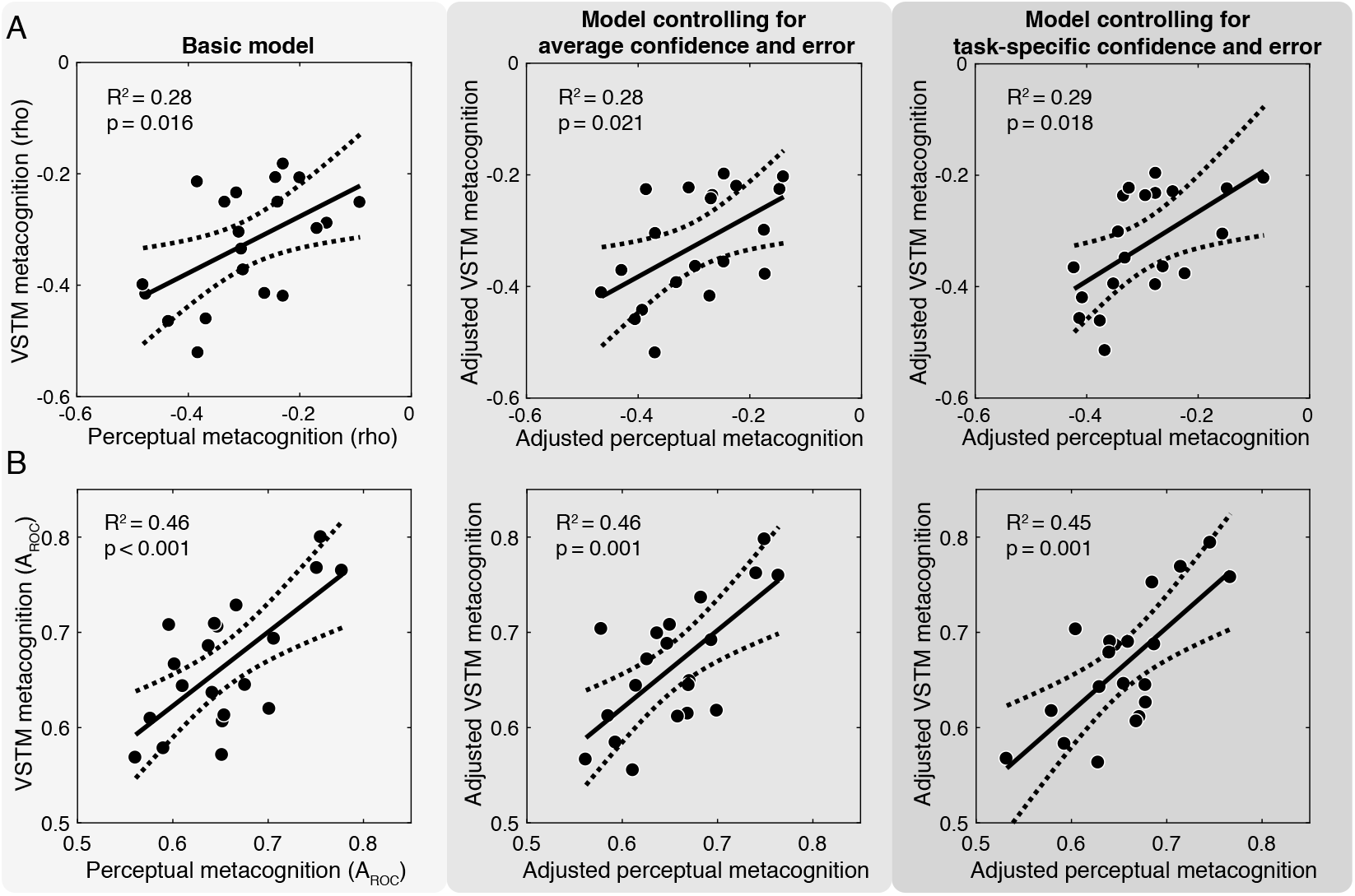
Positive relationship between perceptual and VSTM metacognition in Experiment 1.(A) Cross-task regression using confidence-error correlations as the metric of metacognition.Increasingly complex regression models controlling for task performance and confidence shown from left to right (see Methods). (B) Same models as in A, but using the area under the type 2ROC curve (AROC) as a measure of metacognitive performance. Dashed lines denoted 95%confidence intervals on the linear fit. Black points are individual subjects.

Additionally, we verified that the results of this analysis were robust to our particular metric of metacognition by repeating all analyses using the non-parametric area under the type 2 receiver operating characteristics curve (AROC; (21–23) as our measure of metacognitive accuracy. This measure is obtained by taking the area under the curve formed by plotting the type 2 false alarm rate by the type 2 hit rate at different type 2 criteria. A type 2 false alarm is an incorrect but high confidence trial and type 2 hit is a correct and high confidence trial and the number of confidence criteria is the number of ratings on the scale minus 1. At values of 0.5, this metric indicates that confidence ratings do not discriminate between correct and incorrect trials and values of 1 indicate perfect discriminability. AROC was computed using the method outlined in (21). Because this metric requires binarizing the data into correct and incorrect responses, we defined thresholds for each subject based on the 75^th^ percentile of their response error distributions such that a trial with error larger than this threshold was considered incorrect. This analytically set performance at 75% for each subject, equating accuracy for this analysis. Using a common threshold of 25° for each subject did not change the statistical significance of any analyses reported with this metric. Prior to any analysis, trials with response times below 200 milliseconds or above the 95^th^ percentile of the distribution of response times across all subjects were excluded. The same trial exclusion procedure was applied to both experiments.

### Statistics

We used linear regression to predict individual differences in VSTM metacognition from variation in perceptual metacognition scores (Figures 2 and 4). In a first, “basic model”, we considered only these two variables. Then, to control for individual differences in task performance and confidence ratings, we ran two additional regression models. One included each subject’s mean error and mean confidence as covariates (3 predictors in total) and the other included task-specific confidence and error as covariates (i.e., mean perceptual error and confidence and mean VSTM error and confidence; 5 predictors in total). These three models were run for each metric of metacognition (r values and AROC; see above) and for both experiments. To test for linear effects of confidence on error (Figures 1C and 3D) we regressed single-trial confidence ratings on absolute error for each subject and task and tested the resulting slopes against zero at the group level using a t-test. To test for performance differences between tasks we compared median absolute error between the perception and VSTM task with a paired t-test. We additionally tested for a linear effect of delay period duration in Experiment 2 (Figure 3B) by fitting slopes to each subject’s single-trial absolute error by delay period data and testing these slopes against zero at the group level with a t-test. All tests were two-tailed.

**Figure 3.**
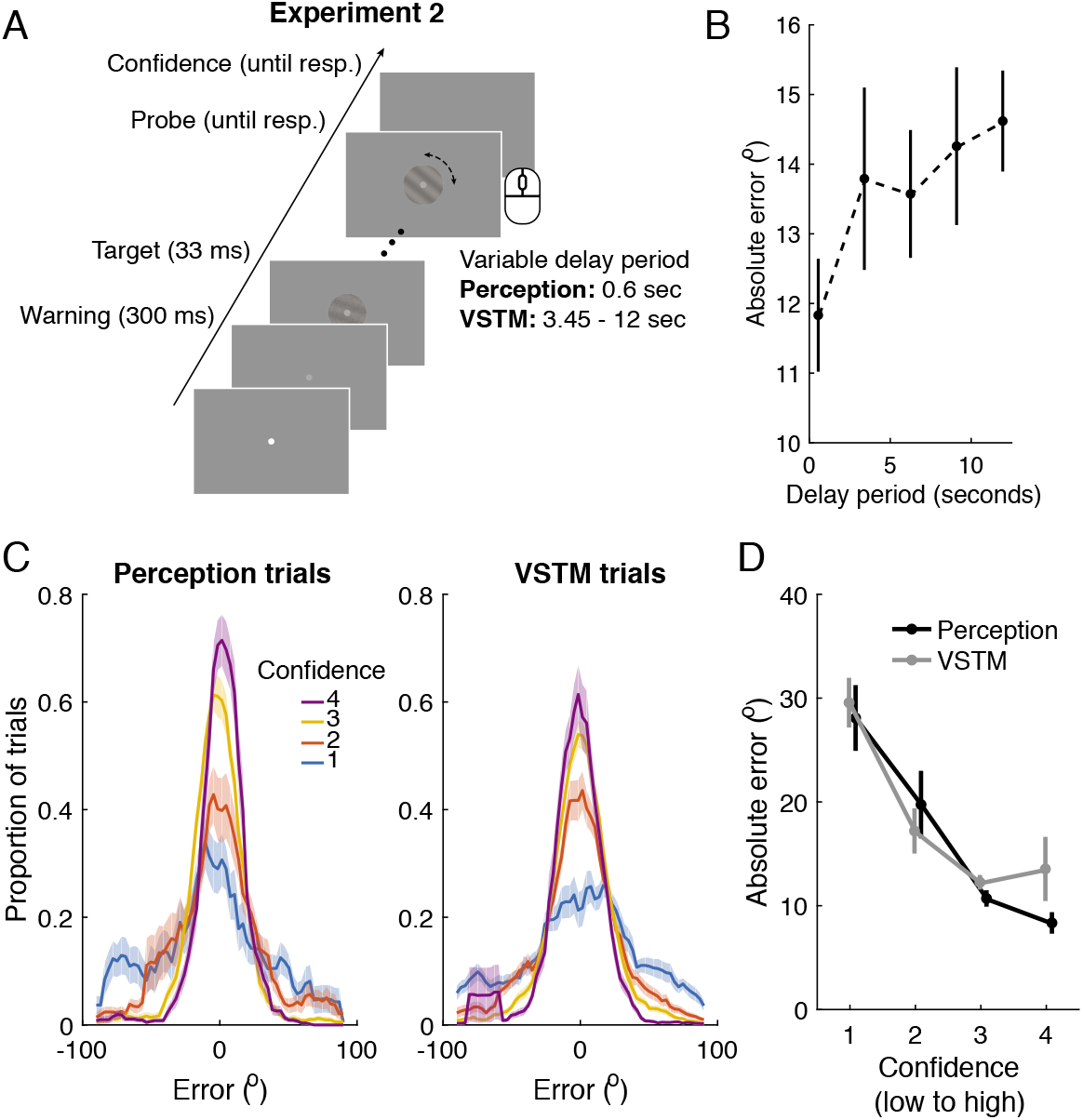
Task and behavior for Experiment 2. (A) Perceptual trials (delay 0.6 seconds) andVSTM trials (delay between 3.45 and 12 seconds) were intermixed within blocks. (B) Error increased with increasing delay length, indicating a loss of information when the orientation needed to be maintained in VSTM. (C) Response error distributions show a clear scaling with confidence. (D) Error decreased as confidence increased in both perceptual and VSTM trials. Shaded bands and error bars indicate ± 1 SEM.

### Experiment 3

This experiment was conducted to test whether the correlation between perceptual and VSTM metacognition depended on both tasks sharing the same task-relevant stimulus feature (e.g. orientation). To this end, we compared metacognition in an orientation perception task and a contrast perception task (Figure 5A) to metacognition in an orientation VSTM task. If the perception of and short-term memory for orientation depend on overlapping representations (13,14), then individual differences in metacognition may be correlated between orientation perception and orientation VSTM, but not between contrast perception and orientation VSTM. Furthermore, we used 2-choice discrimination tasks in Experiment 3 which are amenable to a recently developed Bayesian hierarchical model-based analysis of metacognition that controls for individual differences in task performance and confidence biases while appropriately accounting for variability in individual-subject parameter estimates at the group level (24).

### Participants

Twenty subjects (mean age = 21.8 years, *SD* = 3.18, 13 female) from the University of Wisconsin-Madison community participated in these experiments in exchange for monetary compensation.

### Stimuli

Sinusoidal luminance gratings subtending 2 DVA were centered 1.5 DVA to the right and/or left of fixation (Figure 5A). As in Experiments 1 and 2, the gratings were averaged with white noise. The contrast of the grating and the noise components were adjusted for each subject using a staircase procedure (see below). Stimuli were presented on an iMac computer screen (52 cm wide × 32.5 cm tall; 1920 × 1200 resolution; 60 Hz refresh rate) and viewed by subjects from a chin rest 62 cm away. Stimuli were generated and presented using the MGL toolbox (http://gru.stanford.edu) running in MATLAB 2015b (MathWorks, Natick, MA, USA).

### Contrast perception task

Subjects were instructed to indicate whether the left or right stimulus contained a higher contrast grating using the left and right arrow keys, respectively. Subjects then indicated their confidence using number keys 1-4, where 1 denotes a “complete guess” and 4 denotes “very confident”. Subjects were encouraged to use the full range of the scale and were instructed to understand a number 4 rating as indicating the highest confidence they would feel in this task, given the difficult nature of the task. This confidence rating procedure was the same for all three tasks in this experiment. A target and a standard stimulus were presented simultaneously to the left and right of fixation for 50 ms. The location containing the target was randomly determined on each trial. Each stimulus also had a randomly and independently determined orientation that was task irrelevant. Responses could be made as soon as the stimuli were presented and there was no time limit for responding. The standard stimulus was created by averaging a 10% Michelson contrast grating with an 80% contrast noise patch and the contrast of the target grating was adapted for each subject with a staircase procedure (see below).

### Orientation perception task

This task required subjects to indicate whether the two gratings had the same or different orientation. Both stimuli appeared simultaneously for 50 ms and then subjects indicated their choice followed by their confidence. On “same” trials, both stimuli had an identical orientation, which was randomly determined on each trial (between 1 and 180°), whereas on “different” trials one stimulus was offset by 25° clockwise or counter-clockwise (randomly determined). Whether a trial was same or different was randomly determined. Difficulty was controlled by a staircase procedure that adapted the contrast of both stimuli.

### Orientation VSTM task

This task also required that subjects indicate whether two gratings were the same or different, but here there was a temporal delay of 3 seconds between the first and second stimulus. Thus, the orientation of the first stimulus must be maintained over the delay in order to perform the task. Each grating was presented for 50 ms and subjects provided their choice and confidence, with no time pressure, following the second “probe” stimulus. As in the orientation perception task, both stimuli had an identical randomly determined orientation on same trials, and a difference in orientation of 25° (clockwise or counter-clockwise) on different trials. Trial type was randomly determined as was the location (left or right of fixation) of both stimuli, although the location of both stimuli was always the same for any given trial.

### Staircase

Each task began with 60 trials of a 1-up, 3-down staircase procedure, aimed to converge on ∼80% accuracy. During these 60 trials, stimulus contrast was adjusted according to the PEST algorithm (17), with a starting step size of 8% contrast for all task. For the contrast perception task, the staircase adapted the contrast of the grating component of the target and modulated the contrast of the noise component of the target in the opposite direction so that the overall stimulus contrast always matched the standard (see Figure 5A *left*). For example, if the contrast of the grating component of the target was +12% relative to the grating component of the standard then the noise component of the target was reduced by 12%, thereby matching total stimulus contrast between the target and standard, but producing higher contrast in just the grating component of the target. Starting contrast of the target grating was +20% relative to the standard. For the orientation perception and VSTM tasks the starting contrast of the grating component of each stimulus was 30%, which was averaged with 80% contrast noise. After these initial PEST trials, a threshold was computed as the mean contrast from the last 4 staircase reversals. The staircase continued throughout the duration of each task but with a fixed step size of 0.5% for the contrast task and 0.25% for the orientation tasks, with the starting threshold determined from the initial PEST staircase in the case of the first block of each task, or from the mean of the last 4 reversals from the most recent block.

### Procedure

Each subject completed 3 blocks of 100 trials each for each of the three tasks, resulting in 300 trials per task (with the exception of one subject who only completed 100 trials of the contrast perception task). Blocks of the same task were completed sequentially and task order was randomized. Prior to the start of each new task, subjects completed 60 trials of the initial PEST staircase corresponding to that task. These 60 trials were not included in any analysis. Total task time was ∼1.5 hours.

### Model-based analysis of metacognition

Because Experiment 3 employed 2-choice discrimination tasks we quantified metacognition in a bias-free signal detection theory model (20,24). We used an estimate of metacognitive efficiency (M-ratio) that quantifies the extent to which confidence ratings discriminate between correct and incorrect decisions (i.e., type 2 performance), given the underlying difficulty of the discrimination itself (i.e., type 1 performance), thereby optimally controlling for task difficulty and confidence biases (2). M-ratio is the ratio between the d’ estimated from the confidence data according to a metacognitively ideal observer and the actual d’ computed from task performance. Because both metrics are in the same units, M-ratio will approach 1 if all the information used for the type 1 decision was also available to the type 2 decision, indicating optimal metacognition. Values below 1 reflect suboptimal metacognition.

We used a recently introduced hierarchical modeling approach to estimate the cross-task correlation between individual differences in M-ratio, as is implemented in the freely available toolbox HMeta-d (24, https://github.com/smfleming/HMM) for MATLAB. This toolbox is a hierarchical Bayesian extension of Maniscalco and Lau’s (20) meta-d’ model. The advantage of a Bayesian model in this context is that the estimation of group-level parameters of interest (i.e., M-ratio correlation coefficient across tasks) takes into account parameter uncertainty at the single subject level. This means that a subject whose M-ratio is estimated with high uncertainly will contribute less to the group-level parameter estimate than a subject whose M-ratio is estimated more precisely. In typical maximum likelihood or sum of squares fitting (20), this knowledge of parameter uncertainly is discarded. Simulations suggest this approach produces more accurate parameter recovery and lower false positive rates than non-Bayesian alternatives (24). Cross-task M-ratio correlations were estimated using the HMeta-d function *fit_meta_d_mcmc_groupCorr.m.*

Posterior distributions of parameters were sampled using Markov-Chain Monte-Carlo methods (MCMC) implemented in JAGS (http://mcmc-jags.sourceforge.net). We ran 3 chains of 20,000 samples each with 5,000 burn-in samples. Each subject’s log(M-ratio) for each domain are specified as draws from a bivariate Gaussian. We used a weakly informative normal prior on log(M-Ratio) which encompasses estimates from 167 previous subjects (24), and a uniform prior between −1 and 1 for the correlation coefficient. To assess convergence we ensured that all MCMC chains were well mixed and that the Gelman and Rubins R^ convergence statistics were between 1 and 1.1. Statistical significance for each correlation was assessed by computing the proportion of MCMC samples that fell below zero, multiplied by 2 (akin to a two-tailed non-parametric frequentist test) and by computing 95% high-density intervals (HDI) on the correlation posterior distributions.

## Results

### Experiment 1

Distributions of response error as a function of confidence are shown in Figure 1B. Absolute error significantly decreased with increasing confidence for both the perceptual task (*t*(19) = −13.48, p < 0.0001) and the VSTM task (*t*(19) = −14.88, p < 0.0001), indicating that subject’s confidence reasonably reflected their task performance at the group level (Figure 1C). Error was also significantly greater in the VSTM task as compared to the perceptual task (*t*(19) = −2.10, p = 0.049), reflecting an expected degradation of orientation information when the task required short-term memory maintenance. Confidence ratings were distributed similarly for perception and VSTM tasks (Supplementary Figure 1), and average confidence ratings did not significantly differ between tasks (p = 0.15). See Supplementary Figure 2 for a breakdown of accuracy and response time by block.

Central to our hypothesis, we found a positive relationship across individuals between perceptual metacognition and VSTM metacognition (Figure 2). This relationship was observed when using confidence-error correlations as the measure of metacognition (slope = 0.51, t = 2.63, predictor R^2^ = 0.28, p = 0.016; Figure 2A) and, importantly, was still present after controlling for average confidence and error (slope = 0.54, t = 2.55, predictor R^2^ = 0.28, p = 0.021) and in the model controlling for task-specific confidence and error (slope = 0.62, t = 2.66, predictor R^2^ = 0.29, p = 0.018). All covariate predictors in both control models were not statistically significant (ps > 0.30). These results indicate that, although the confidence-error correlation may be influenced by task performance and confidence biases, these factors did not account for the across-subjects correlation between perceptual and VSTM metacognition.

The same relationship was observed when using AROC as the metric of metacognition (Figure 2B). With the basic model, perceptual metacognition significantly predicted VSTM metacognition (slope = 0.77, t = 3.96, predictor R^2^ = 0.46, p = 0.0009). This relationship held when controlling for average confidence and error (slope = 0.82, t = 3.78, predictor R^2^ = 0.46, p = 0.0016) and when controlling for task-specific confidence and error (slope = 0.88, t = 3.85, predictor R^2^ = 0.45, p = 0.0017). As before, all other covariate predictors across both control models were non-significant (ps > 0.26). We examined the correlation between all predictor variables in all of our models (Table 1) and found that there was collinearity between several, quite expected, covariate predictors (e.g., average confidence predicted average error, perceptua confidence predicted VSTM confidence). Importantly, however, there were no significant correlations between our predictors of interest (both perceptual metacognitive scores, AROC or rho) and any other covariate predictors, indicating that task performance and confidence are unlikely to be driving the cross-task correlation in metacognition. These results indicate that the relationship observed between perceptual and VSTM metacognition was independent of the particular metric used and was not accounted for by correlated individual differences in task performance or average confidence.

**Table 1.** Correlation matrix for every predictor in each model in Experiment 1 and Experiment 2(gray regions). X’s denote predictor combinations that were not used in any model. Significant correlations (p<0.05) are noted in bold and with asterisks.

| Predictor – Exp. 1, Exp. 2 | 1 | 2 | 3 | 4 | 5 | 6 | 7 | 8 |
| --- | --- | --- | --- | --- | --- | --- | --- | --- |
| 1. Perceptual metacognition (rho) | - | -0.13 | -0.03 | -0.16 | -0.04 | -0.10 | -0.01 | X |
| 2. Average confidence | 0.33 | - | -0.41 | X | X | X | X | 0.15 |
| 3. Average error | 0.01 | **-0.54\*** | - | X | X | X | X | 0.01 |
| 4. VSTM confidence (average) | 0.33 | X | X | - | -0.42 | **0.95\*** | -0.38 | 0.17 |
| 5. VSTM error (average) | 0.08 | X | X | **-0.54\*** | - | -0.42 | **0.91\*** | -0.01 |
| 6. Perceptual confidence (average) | 0.32 | X | X | **0.94\*** | **-0.44\*** | - | -0.38 | 0.13 |
| 7. Perceptual error (average) | -0.05 | X | X | **-0.53\*** | **0.83\*** | **-0.53\*** | - | 0.03 |
| 8. Perceptual metacognition ($A_{ROC}$) | X | -0.29 | 0.07 | -0.28 | 0.01 | -0.14 | -0.29 | - |

### Experiment 2

This experiment served to replicate the cross-task correlation observed in Experiment 1 while further minimizing procedural differences between tasks by intermixing perceptual and VSTM trials of differing delays (Figure 3A). Error increased monotonically with delay duration (*t*(19) = 2.85, p = 0.010. Figure 3B), and perception trials had lower error than VSTM trials, collapsing across delays (*t*(19) = 3.33, p = 0.003), indicating the expected loss of information in VSTM relative to perception. As in Experiment 1, error decreased with increasing confidence during both perception (*t*(19) = −7.56, p < 0.0001) and VSTM trials (*t*(19) = −8.99, p < 0.0001), indicating that confidence reliably tracked performance at the group level (Figure 3C & 3D). Average confidence was lower on VSTM (mean = 2.59, SEM = 0.133) than on perception trials (mean = 2.71, SEM = 0.132; *t*(19) = 2.97, p = 0.0077; Supplementary Figure 1).

Importantly, we replicated the positive relationship between perceptual and VSTM metacognition with quantitatively better model fits in a new set of subjects. Using confidence-error correlations (Figure 4A) perceptual metacognition robustly predicted VSTM metacognition in the one-predictor basic model (slope = 0.60, t = 5.21 predictor R^2^ = 0.60, p < 0.0001), the three-predictor model controlling for average confidence and error (slope = 0.60, t = 4.91, predictor R^2^ = 0.59, p = 0.0001), and in the five-predictor model controlling for task-specific confidence and error (slope = 0.59, t = 4.44, predictor R^2^ = 0.58, p = 0.0005. All covariate predictors in both control models were non-significant (ps > 0.64). As in Experiment 1, some covariate predictors were significantly correlated (Table 1; gray region), but no covariates were significantly correlated with perceptual metacognition, the predictor of interest. This effect was also observed when using AROC as the metric of metacognition for the basic model (slope = 0.53, t = 3.88, predictor R^2^ = 0.45, p = 0.0011), the three-predictor model (slope = 0.52, t = 3.58, predictor R^2^ = 0.44, p = 0.002), and the five-predictor model (slope = 0.54, t = 3.43, predictor R^2^ = 0.44, p = 0.004). All covariates in both control models were non-significant (ps > 0.52).

**Figure 4.**
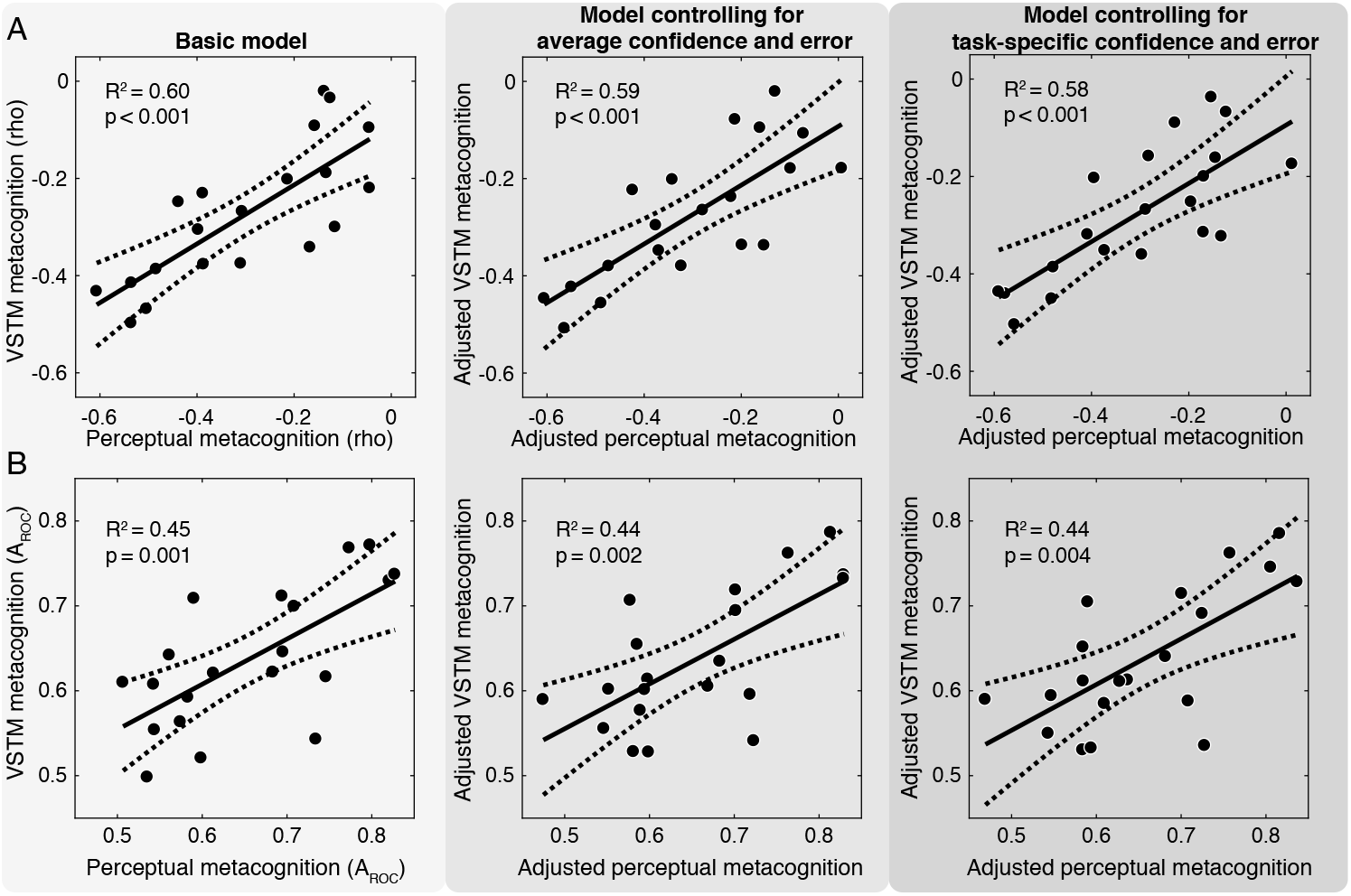
Replication of the positive relationship between perceptual and VSTM metacognition in Experiment 2. (A) Same regression models as in Figure 2, indicating the cross-task relationship using confidence-error correlations as the metric of metacognition. (B) Same as inA, but with AROC as the metric of metacognition. Dashed lines denoted 95% confidence intervals on the linear fit.

### Experiment 3

This experiment was conducted to test whether correlated individual differences in metacognition depended on the perception and VSTM tasks sharing the same task-relevant stimulus feature (i.e., orientation). Task accuracy (% correct) and metacognitive efficiency (M-ratio) for each task and subject are shown in Figure 5B. Accuracy was comparable between the contrast perception task (mean = 78.6%, SEM = 0.012) an the orientation perception task (mean = 80.4 %, SEM = 0.009; p = 0.33), as well as between the contrast perception task and the orientation VSTM task (mean = 76.5%, SEM = 0.018%, p = 0.39), but differed significantly between the orientation perception and the orientation VSTM tasks (*t*(19) = 2.68, p = 0.015). This is party because four subjects reached the maximum contrast allowable by the staircase (25%), so the VSTM task never got easier for them. In general, higher target contrasts were needed in the VSTM task (mean = 15.5%, SEM = 0.012) compared to both the orientation perception task (mean = 8.5%, SEM = 0.005; *t*(19) = 6.50, p < 0.0001), and the contrast perception task (mean = 6.2%, SEM = 0.005; *t*(19) = 6.49, p < 0.0001), indicating that more signal was needed in the VSTM task to achieve threshold performance. Metacognitive efficiency, on the other hand, did not significantly differ between tasks (ps > 0.16; see Supplementary Figure 3 for full posterior distributions of M-ratio for each task, and comparisons between tasks), and was well below the optimal M-ratio of 1 for all tasks (contrast perception: mean M-ratio = 0.4, HDI = [0.27, 0.54], orientation perception: mean = 0.44, HDI = [0.30, 0.59], orientation VSTM: mean = 0.53, HDI = [0.38, 0.68]). Mean confidence (see Supplementary Figure 1) also did not differ between either of the perceptual tasks and the VSTM task (ps > 0.08), but was significantly lower in the contrast perception task as compared to the orientation perception task (*t*(19) = 3.06, p = 0.006).

To our primary question of correlated individual differences in metacognition, we observed a large positive correlation between metacognitive efficiency in the orientation perception task and orientation VSTM task (Figure 5C; rho = 0.90, HDI = [0.58 0.99], p < 0.0001), consistent with the results of Experiments 1 and 2. However, we did not observe a significant correlation between metacognition in the contrast perception task and the orientation VSTM task (rho = 0.32, HDI = [-0.32 0.85], p = 0.32), suggesting that the correlation between perception and VSTM metacognition depends on both tasks sharing the same task-relevant stimulus feature. Interestingly, metacognition was also not correlated between the two perceptual tasks (rho = 0.24, HDI = [-0.35 0.76], p = 042), again underscoring the importance of the similarity of stimulus feature used. We also computed the difference between these correlation distributions and found that the correlation between orientation perception and orientation VSTM memory was significantly larger than the correlation between contrast perception and orientation perception (HDI = [0.07 1.27], p = 0.027) and trending larger than the correlation between contrast perception and orientation VSTM (HDI = [-0.03 1.24], p = 0.063).

**Figure 5.**
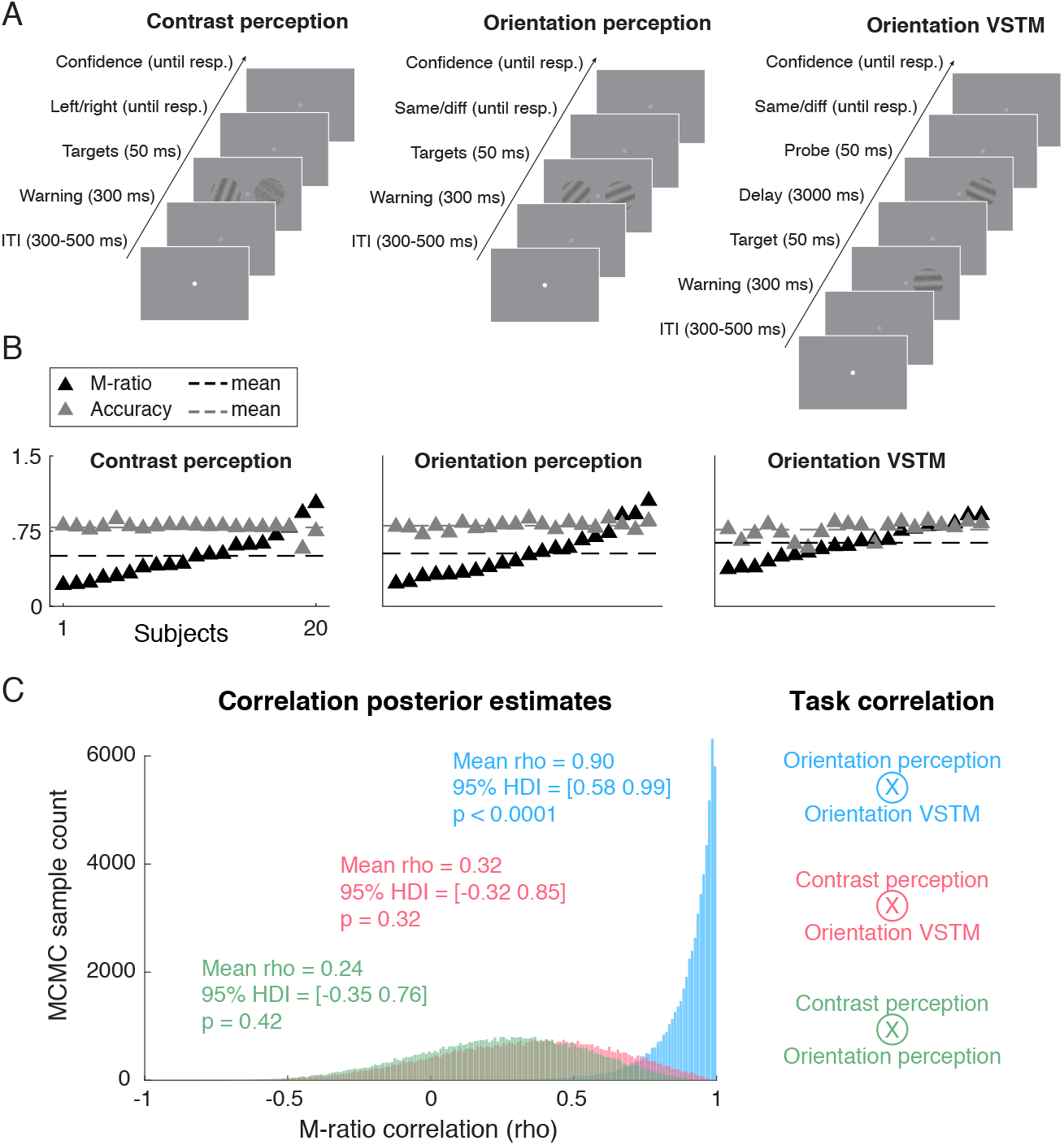
Tasks, behavior, and metacognitive correlations from Experiment 3. (A) To compare metacognition across discrimination tasks while varying task and the task-relevant stimulus feature, subjects performed 1) a contrast perception task, judging which stimulus contained ahigher contrast grating, 2) an orientation perception task, judging whether the two stimuli had the same orientation, and 3) an orientation VSTM task, judging whether a memorized target grating had the same orientation as a probe grating that appeared 3 seconds later. (B)Individual subject accuracies (proportion correct) and estimates of metacognitive efficiency (M-ratio) for each task. Note that because M-ratio for each subject is estimated in the same model,estimates are not fully independent (24). (C) Posterior distributions of cross-task correlations in metacognitive efficiency, which reveal a strong positive correlation between orientation perception and orientation VSTM metacognition, but not between other tasks.

## Discussion

Metacognition is an important aspect of decision-making (25,26), learning (27), development (28), and perhaps certain aspects of conscious experience (29,30), and can be compromised in psychiatric disorders (31–33). It is currently unclear whether an individual with good metacognitive ability in one domain also has good metacognition in other domains. In Experiment 1, we found that individuals with more accurate metacognition in perceptual judgments also showed more accurate metacognition in a VSTM task requiring stimulus maintenance over a 7 second delay period. This relationship was present when using two different measures of metacognitive performance and regression models controlling for task performance and mean confidence revealed that this effect was not driven by correlated individual differences in task performance or confidence biases. We then replicated these findings in Experiment 2 with a new set of subjects using a task that intermixed perceptual and VSTM trial types within blocks. Intermixing trial types in Experiment 2 more than doubled the proportion of variance in VSTM metacognition explained by perceptual metacognition in the models using error-confidence correlations relative to Experiment 1 when trial types were blocked (mean increase in R^2^ = 0.30, a factor of ∼2.2), highlighting the importance of minimizing procedural differences between tasks. A comparable increase across experiments was not seen, however, when using the AUC metric, which already showed a very large effect size in both experiments and across all models (mean R^2^ = 0.45, Cohen’s *d* = 1.81). In Experiment 3 we compared an orientation VSTM task to an orientation perceptual task and a contrast perception task. We again found a large positive correlation (R^2^ = 0.81) between metacognition in the orientation VSTM and orientation perception task, but not between the orientation VSTM and contrast perception task, nor between contrast perception and orientation perception tasks, highlighting the importance of both tasks sharing the same relevant stimulus feature. Importantly, given known biases in VSTM metacognitive judgments (34), metacognition in Experiment 3 was quantified within a signal detection theory model (20,24) that controls for confidence biases and task performance. Taken together, these results provide the first evidence in humans for a medium-to-high positive correlation between an individual’s metacognitive abilities in perception and VSTM, when both domains share a common stimulus feature.

The present results contrast with recent experiments examining the relationship between metacognition of visual perception and long-term memory, which have typically observed no correlation (4-6; but see 7). We reason that, in contrast to long-term memory, VSTM for a given stimulus feature is thought to rely on the same neural representations that support perception of that stimulus feature (11–14), and this may underlie the cross-task correlation in metacognitive performance. This explanation follows naturally from “first-order” models of metacognition according to which confidence and task performance are driven by the same internal representation of stimulus evidence (35–38). For example, in signal detection theoretic models, the absolute distance of the decision variable from the decision criterion is a proxy for confidence (39,40). Thus, if perception and VSTM were supported by the same internal representation of the stimulus, then the computation of confidence across the two tasks would also be based on the same representations, leading to correlated behavior. “Second-order” models of metacognition, in contrast, posit an architecture with a secondary confidence read-out process, which may be influenced by additional sources of noise (41) or other signals not directly related to the stimulus, such as action-related states (42,43), cortical excitability (44), or arousal (45,46).

The results of Experiments 1 and 2 are also compatible with second-order models of metacognition, although several possible relationships between first- and second-order processes could explain our findings. Shared first-order (sensory) representations across tasks might be enough to produce a behavioral correlation despite separate second-order readout mechanisms. Alternatively, both first- and second-order processes may be shared across tasks, or only the second-order process shared, though this latter possibility is unlikely given existing neural evidence for shared representations in visual regions across perception and VSTM (14,47,48). The results of Experiment 3, however, provide support for 1st-order models because they suggest that shared sensory representations underlie the cross-task correlation in metacognition. Because metacognition was not reliably correlated when tasks differed in their relevant stimulus feature, even when both tasks were perceptual, this points towards a first-order model of metacognition. Yet another alternative is that the correlation was dependent on the task structure, for example, because both orientation tasks involved same/different judgments. This account may also explain why a previous report comparing metacognition for contrast and orientation judgments in the context of a visual search paradigm did find correlated individual differences (8), but recent work comparing a variety of perceptual paradigms with different task structures and stimuli did not find a correlation (49).

Although the present findings are consistent with a domain-general model of metacognition for perception and VSTM, correlations at the behavioral level raise further questions about what specific aspects of metacognitive processing are shared. For example if one’s ability to learn stable confidence criteria over time improves metacognitive accuracy (38), then metacognitive abilities may be high across domains for an individual with superior learning abilities, perhaps related to recent work implicating hippocampal myelination in perceptual metacognition (50). However, this need not imply that the underlying neural substrate responsible for computing the appropriate levels of confidence is itself domain-general. Similarly, recent work has highlighted specific factors beyond stimulus evidence that modulate confidence, leading to dissociations of confidence and task performance within an individual (15,51,52). For example, spontaneous trial-to-trial fluctuations in oscillatory neural activity in the alpha-band (8-13 Hz), which are thought to reflect visual cortical excitability (53,54), have been shown to bias confidence ratings, but not objective performance in a visual discrimination task (44). Perhaps a subject who is less susceptible to such influences from sources not directly related to the difficulty of stimulus discrimination would show better metacognition across different domains. Future work examining neural correlates of metacognitive performance across different domains may contribute in a substantive way to this issue. As an example, McCurdy and colleagues (7) observed a positive correlation between metacognition of perception and recollection memory at the behavioral level, but found distinct (as well as overlapping) neural structures whose gray matter volume related to metacognitive performance in the different tasks. This suggests that only a portion of the processing stages or computations involved in generating confidence need be shared across tasks in order to produce a behavioral correlation. Nevertheless, the experiments reported here provide an important first step for future work by demonstrating a clear correlation between metacognitive behavior in perception and VSTM

## Acknowledgments

We would like to thank John B. Barrett, Missy Switzky, and SawyerCimaroli for help with data collection and two anonymous reviewers for their helpful feedback. Supported by MH095984 to B.R.P

